# In vitro reconstituted quorum sensing pathways enable rapid evaluation of quorum sensing inhibitors

**DOI:** 10.1101/2021.10.29.466404

**Authors:** Dingchen Yu

**Author notes:** Corresponding author: Dingchen Yu.

## Abstract

Quorum sensing, as inner- or inter-species microbial communication process orchestrated by diffusible autoinducers, typically results in collective pathogenic behaviours, being recognized as a promising druggable target for anti-virulence. Here, we reconstituted *las* and *rhl* quorum sensing pathways of *Pseudomonas aeruginosa*, mediated by acyl-homoserine lactones (AHLs) and LuxI/LuxR-family proteins, with fluorescence output in *Escherichia coli* cell-free expression system, offering a platform to rapidly evaluate quorum sensing inhibitors (QSIs) *in vitro*. Previously reported small-molecule quorum sensing inhibitors for interfering with *P. aeruginosa* quorum sensing systems were tested and showed mild to high on-target inhibition as well as off-target toxicity. Of note, quercetin displayed potent on-target inhibition to quorum sensing pathways as well as acceptable off-target toxicity to cell-free expression machinery. Upon our work, cell-free platform is anticipated to further facilitate automated and high-throughput drug screening, bridge *in silico* and *in vivo* drug-screening methods, and accelerate the upgrading of antimicrobial arsenal.

**GRAPHICAL ABSTRACT:** 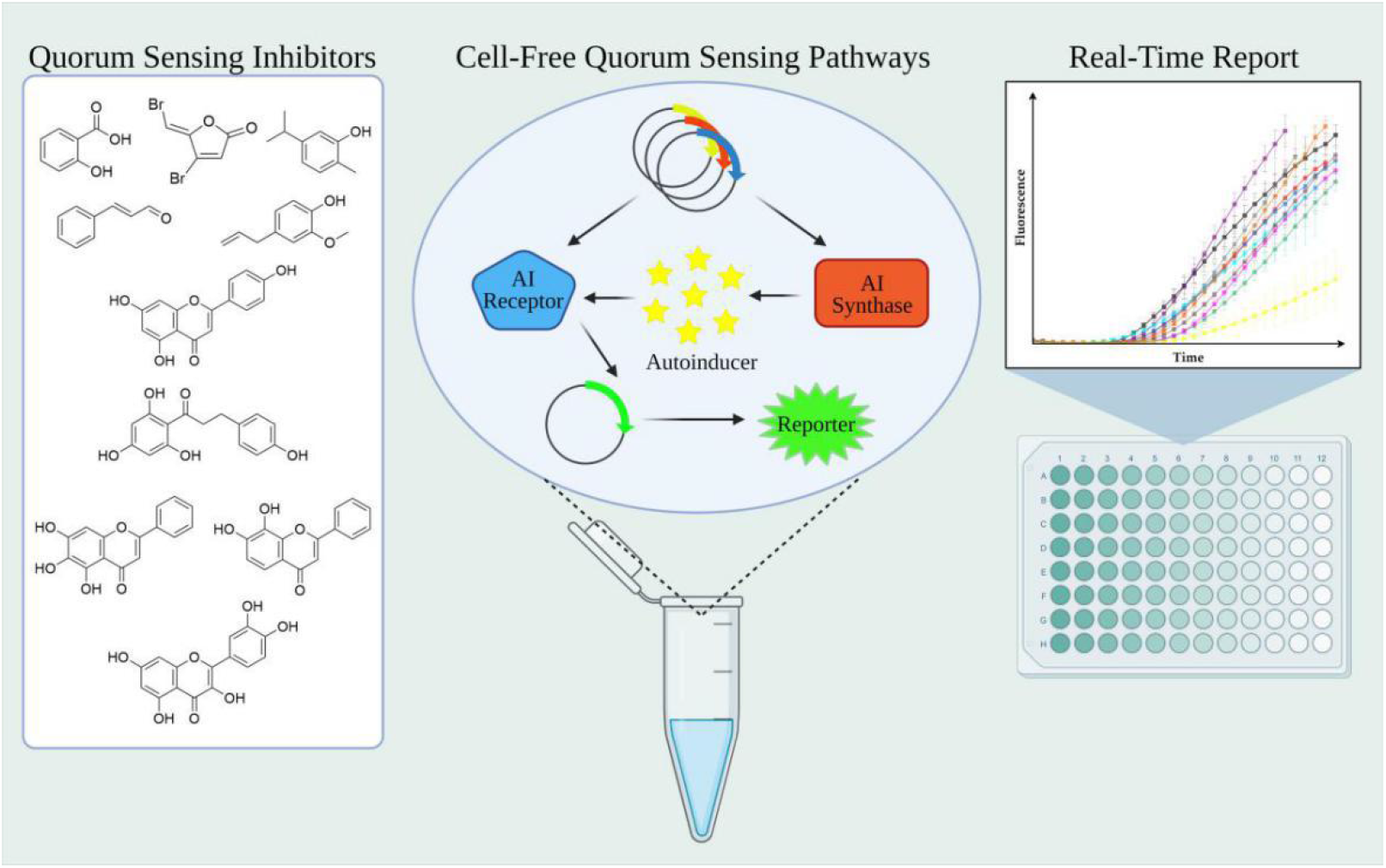

## INTRODUCTION

Quorum sensing (QS) is inner- or inter-species communication process widespread in microbial world, which enables microorganisms to coordinate gene expression by diffusible autoinducers (AIs), and to make decisions synchronously in a cell density-dependent manner, such as bioluminescence, biofilm formation, and virulence factor excretion [1]. Bacterial quorum sensing systems can be grouped based on the type of autoinducers, including acyl-homoserine lactones (AHLs) in Gram-negative bacteria, autoinducer peptides (AIPs) in Gram-positive bacteria, AI-2 shared among diverse species, and other compounds [2]. The discovery of quorum sensing could date back to 1970s when a marine bacterium, *Aliivibrio fischeri*, was reported to express bioluminescence only after its population density reaches a threshold [3]. The first quorum sensing system, *lux* pathway, was later characterized, in which AHL-type AI molecule N-(3-oxo-hexanoyl)-homoserine lactone (3-oxo-C6-HSL) is synthesized by LuxI and sensed by allosteric transcription factor LuxR to drive expression of luciferase operon *luxCDABE* for bioluminescence generation [4]. As a representative of quorum sensing systems, the *lux*-type pathways have been extensively discovered in *Vibrio cholerae, Vibrio harveyi, Pseudomonas aeruginosa, Agrobacterium tumefaciens* and numerous other Gram-negative bacteria [5], and different systems usually utilize AHL signals with varied acyl chain length and/or modification.

Ecologically, quorum sensing endows the host species with communication tools to respond to competitors in microbiota or harsh environmental conditions as a collective, in order to compete for a niche. Several pathogenic bacteria, like *Pseudomonas aeruginosa*, harness quorum sensing as a pivotal means to control virulence expression [6]. Instead of expressing virulence-related genes sporadically, pathogenic bacteria utilized QS-mediated synchronization to maximize the profits of expressing resource-intensive genes, including efflux pumps for antibiotic resistance, exotoxins for cytotoxicity, proteases for immune evasion, and rhamnolipids for biofilm construction [6], consequently creating more chances for colonization.

Because of its crucial part in virulence expression, quorum sensing is being re-considered as a druggable process, and researchers aim to discover or design quorum sensing inhibitors (QSIs) targeting quorum sensing components to interfere with microbial pathogenicity [7,8]. Encouragingly, it is reported that knockout of quorum sensing components or addition of quorum sensing inhibitors renders *P. aeruginosa* more susceptible to an aminoglycoside antibiotic, tobramycin, and its biofilm structure more vulnerable [9-12]. A large amount of quorum sensing inhibitors have been reported over decades, including small molecules, peptides and quorum quenching enzymes, with target QS processes ranging from signal synthesis (along with precursor synthesis), signal transport, signal diffusion, receptor turnover to signal-receptor complex formation [7]. As for AHL-based QS systems, LuxI- and LuxR-family proteins are recognized as hotspots for drug screening. Methodologically, to identify possible small-molecule QSIs from a large library of compounds, researchers often prioritize virtual screening to predict ligand-receptor interactions in bulk, following *in vivo* screening using compounds selected *in silico* [13,45]. The reduction of QS-relevant gene expression usually serves as a gold standard to evaluate the effectiveness of QSIs [9,12]. Yet, considering the complicated metabolic network of pathogenic bacteria, there might exist false-positive “treater” compounds that interfere with non-relevant targets but cause downregulation of several QS-related genes expression (or steady-state concentration of their products), likely diverting researchers to misjudgment. Therefore, instead of experimenting pathogenic bacteria, genetically amenable model bacteria with clearer metabolism background, like *Escherichia coli*, could be engineered, with the introduction of heterologous quorum sensing pathways, as reporter strains to indicate potential quorum sensing inhibition with colorimetric (*e*.*g*., coupled with violacein-producing biosensor bacteria, *Chromobacterium violaceum* CV026 [15]) or fluorescent outcome [12]. Nevertheless, false-positive results are still unavoidable, since any chemicals influencing the vitality of living cells might engender fluctuating gene expression.

Notably, cell-free gene expression (CFE) technology provides synthetic biologists with an alternative option to engineer gene circuits, while circumventing the use of intact cells [16-18]. Comprised of cell lysates and exogenously added components, this bottom-up platform enables rapid reconstitution of complex transcription-translation (TXTL) process and biochemical reactions *in vitro*, expediting applications encompassing point-of-care biosensing [19-22], on-demand therapeutics production [23], metabolic pathway prototyping [24-26], high-throughput enzyme characterization [27] and parts mining [28]. Importantly, CFE platform gets rid of host genome and concomitant unpredictable influences, a meritorious feature that helps scientists develop platforms to characterize noncanonical CRISPR RNAs (ncrRNA) [29] and systematically screen anti-CRISPRs (Acrs) [30]. Inspired by CFE’s unique advantages, we propose to extract quorum sensing pathways from pathogenic bacteria and reconstitute them in a cell-free platform, whereby enabling accurate, *in vitro* evaluation of the inhibitory effects of QSIs, to a great extent decoupling the disturbances of those frequently-used “black-box” systems, namely living cells, with basic needs for survival and growth.

In this work, we set out to reconstruct two *lux*-type quorum sensing pathways of *Pseudomonas aeruginosa, las* and *rhl* pathways, in *Escherichia coli* cell-free expression system. To prototype *las* and *rhl* pathways, cell-free reactions were performed to synthesize LuxR-family allosteric transcription factors (LasR and RhlR) and LuxI-family AHL synthases (LasI and RhlI), and green fluorescent protein (GFP) expression was controlled by corresponding QS promoters (pLas and pRhl) as reporter in both pathways. The successful reconstitution was demonstrated by gauging real-time fluorescence kinetics and detecting AHL biosynthesis by high-performance liquid chromatography-mass spectrometry (LC-MS). As a proof of concept, ten small-molecule quorum sensing inhibitors were tested within hours, exhibiting mild to high on-target inhibition as well as off-target toxicity. Based on these results, our method shows great potential in the rapid evaluation of quorum sensing inhibitors, and we reason that other targeted antimicrobial agents can be evaluated similarly, as the approach of *in vitro* reconstitution proposed herein is applicable to other druggable microbial pathways.

## RESULTS AND DISCUSSION

### Reconstitution of *las* and *rhl* quorum sensing pathways

In a typical *lux*-type quorum sensing process, LuxI-family AHL synthase catalyzes acylation and lactonization of acyl-acyl carrier protein (ACP) and S-adenosyl-methionine (SAM) to produce AHL [31], which diffuses freely through cell membranes and over environment; the cognate LuxR-family transcription factor binds AHL and dimerizes to form active complex, searches for QS-controlled promoters, and recruits RNA polymerases to initiate downstream gene transcription (Figure 1B). *Pseudomonas aeruginosa*, as a notorious human pathogen, possesses two AHL-mediated quorum sensing systems, namely *las* and *rhl* [6]. In the *las* system, LuxI-family AHL synthase LasI produces N-(3-oxo-dodecanoyl)-homoserine lactone (3-oxo-C12-HSL), which is sensed by LuxR-family transcription factor LasR. Similarly, in the *rhl* system, the signal molecule N-butyryl-homoserine lactone (C4-HSL) is produced by RhlI and sensed by RhlR. Owing to the clinical significance of *P. aeruginosa*, there are cell-free biosensors [32] and engineered “sense-and-kill” bacteria [33,34] based on its QS pathways reported, using 3-oxo-C12-HSL or C4-HSL as biomarkers. To our knowledge, though researchers reconstituted complete cell-free *lux* pathway [61], no complete *P. aeruginosa* QS pathway has been reconstituted *in vitro* before.

**Figure 1:**
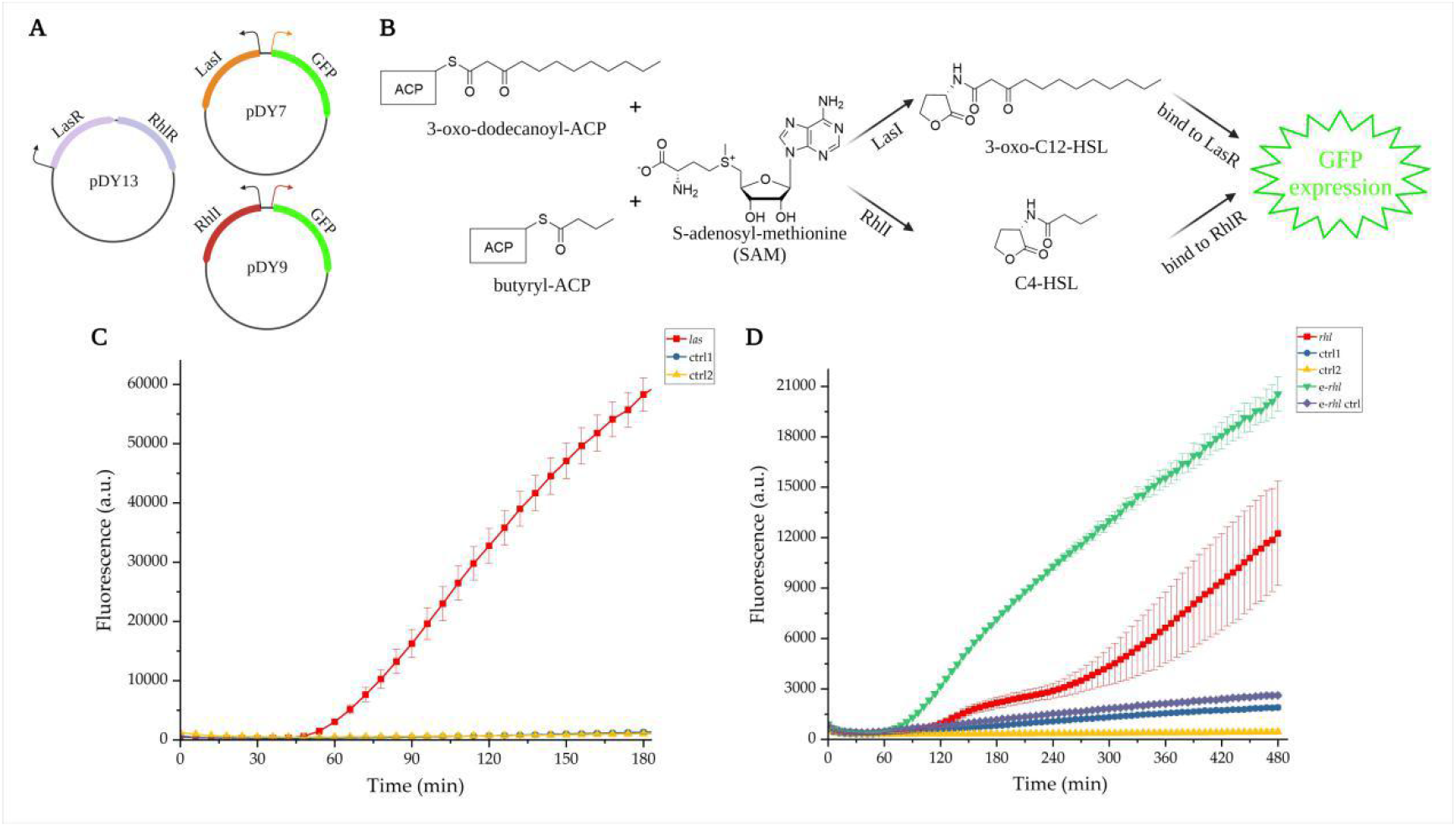
Cell-free *las* and *rhl* pathways. (A) Plasmids used in the reconstitution of *las* and *rhl* pathways, in which black arrow denotes constitutive promoter BBa_J23100; orange arrow denotes *las*-induced promoter pLas; red arrow denotes *rhl*-induced promoter pRhl; GFP denotes green fluorescence protein. (B) Biochemical processes recapitulated in the cell-free *las* and *rhl* pathways, in which ACP denotes acyl carrier protein; 3-oxo-C12-HSL denotes N-(3-oxo-dodecanoyl)-homoserine lactone; C4-HSL denotes N-butyryl-homoserine lactone. (C) 3-hour fluorescence kinetics of *las* pathway (8 nM pDY7, 4 nM pDY13). (D) 8-hour fluorescence kinetics of *rhl* pathway (8 nM pDY9, 4 nM pDY13). The red curves denote *las* and *rhl* pathways; the blue curves (ctrl1) denote control reactions lacking AHL synthases; the yellow curves (ctrl2) denote control reactions lacking transcription factors (8 nM pDY7/9); the green curve (e-*rhl*) denotes enhanced *rhl* pathway and the navy curve denotes its according control reactions lacking AHL synthases (4 nM pDY13, 8 nM pDY10/11). Experimental data generated with microplate reader are presented as mean (s.d.) from sample sizes of n=3 biological replicates drawn from a single batch of *E. coli* extract.

In our earlier attempts of employing *Escherichia coli* cell-free TXTL techniques [25,35], T7 RNA polymerases were harnessed for *in vitro* transcription. Unfortunately, our previously-used protocol [25,35,36] showed poor performance in expressing reporter proteins by σ_70_-driven promoters, but maintained robust expression efficiency using T7 expression system (Figure S2). This indicates activation of *E. coli* transcription machinery is a pre-requisite to *in vitro* pathway prototyping, since the reconstituted QS system will be based on transcription factors (*E. coli* native factors, such as σ_70_, and heterologous QS factors, LasR and RhlR) and bacterial core RNA polymerases instead of T7 RNAP. According to successful examples [28,32,37-40], we discovered that setting appropriate incubation process after lysis, namely ribosomal run-off, greatly enhanced the performance of native expression system, yielding GFP at >300 μg/mL (Figure S2). Dialysis is not included in our improved protocol, since AHL synthases use SAM and acyl-ACP in the lysates as substrate while dialysis would remove these endogenous components.

Upon the optimized CFE platform, we set out to prototype *las* and *rhl* pathways. For each pathway, we co-expressed two plasmids in a cell-free reaction (Figure 1A): one plasmid (pDY13) constitutively expresses two allosteric transcription factors, LasR and RhlR; the other plasmid (pDY7 or pDY9) constitutively expresses AHL synthase, LasI or RhlI, and conditionally expresses GFP by corresponding QS promoters, pLas or pRhl. Also, we set control reactions for each pathway, in which either the first plasmid was not included or the latter plasmid was substituted with a plasmid removing the AHL synthase part while remaining the QS promoter-controlled GFP expression cassette (pDY10 and pDY11, see Supplementary Material). The fluorescence kinetics of reconstituted QS pathways were monitored for hours at 30 °Cwith a microplate reader. As expected, *las* and *rhl* pathways both successfully produced GFP (Figure 1C & 1D, red curves). Additionally, we observed basal expression of reporter proteins in the control reactions lacking QS transcription factors or AHL synthases, indicating low background expression of pLas and pRhl in *E. coli* cell-free system (Figure 1C & 1D, blue and yellow curves). Generally, the reconstituted *las* pathway yielded significantly higher response than the *rhl* pathway, for which we speculate: (i) the strength of LasR-pLas pair to initiate transcription is higher than RhlR-pRhl; (ii) the instability of RhlR renders lagging signal transduction [14]; and (iii) the substrates of *las* pathway are more abundant than *rhl* pathway, as medium- and long-chain acyl-ACP is predominant in *E. coli* [41]. Attempting to enhance the performance of *rhl* pathway, we designed a two-step cell-free reaction. 2.5 μL CFE solution pre-expressing transcription factors for 8 hours was supplemented to the cell-free reaction encoding *rhl* pathway, and the performance of *rhl* pathway improved (Figure 1D, green curve), so as the background of its control reaction (Figure 1D, navy curve).

To further corroborate the successful reconstitution of *las* and *rhl* pathways, we separately analyzed the cell-free reactions with LC-MS. The biosynthesis of 3-oxo-C12-HSL was successfully detected (Figure S3). However, C4-HSL remained covered by the high ionic chemical background noise (Figure S4), which is plausible considering previously-observed limited activity of cell-free *rhl* pathway. So far, we have reconstituted two AHL-mediated *P. aeruginosa* QS pathways *in vitro*, encompassing the basic biochemical processes of signaling, sensing and functioning, laying the foundation for subsequent rapid evaluation of quorum sensing inhibitors.

### Rapid evaluation of quorum sensing inhibitors

Plants are huge reservoirs of quorum sensing inhibitors, and researchers hypothesize that plants accumulate “weapons” (*i*.*e*., secondary metabolites) through evolution to impede and combat diverse endophyte or ectophyte pathogens [14,42]. As for *P. aeruginosa*, a pioneering work reported that synthetic analogues of an alga-derived halogenated furanone inhibited QS-controlled gene expression [9,43]. A plethora of phytochemicals were later reported as potent QSIs to reduce *P. aeruginosa* virulence, including salicylates [13,44], monoterpenoids [47], flavonoids [14,45] and phenylpropanoids [15,44,46]. Indeed, some studies declared QSIs that they discovered target components of *las* or *rhl* pathways, LasI/RhlI or LasR/RhlR, specifically, and demonstrated that typically via molecular docking or reporter bacteria assays. However, the results of molecular docking require further wetware proof. Given living bacteria’s intricate metabolism and uncharacterized regulatory networks, there also lacks a reliable method to interrogate whether select compound interferes with target pathways specifically. Considering the unique bottom-up modality of cell-free expression technology, we commenced to re-evaluate 10 previously-reported QSIs (Figure 2A), including furanone C-30 [9], salicylic acid [13,44], cinnamaldehyde [15,44], eugenol [46], carvacrol [47], and five flavonoid compounds [14,45] (baicalein, quercetin, apigenin, phloretin and 7,8-dihydroxylflavone), based on the *in vitro* reconstituted QS pathways. Of ten molecules, furanone C-30, salicylic acid, eugenol, and the flavonoids are believed to impact LuxR-type transcription factors [9,13,14,45,46]; carvacrol is reported to reduce AHL synthesis [47]; and cinnamaldehyde is predicted to interact with the substrate-binding pocket of LasI for inhibiting 3-oxo-C12-HSL biosynthesis [15].

**Figure 2:**
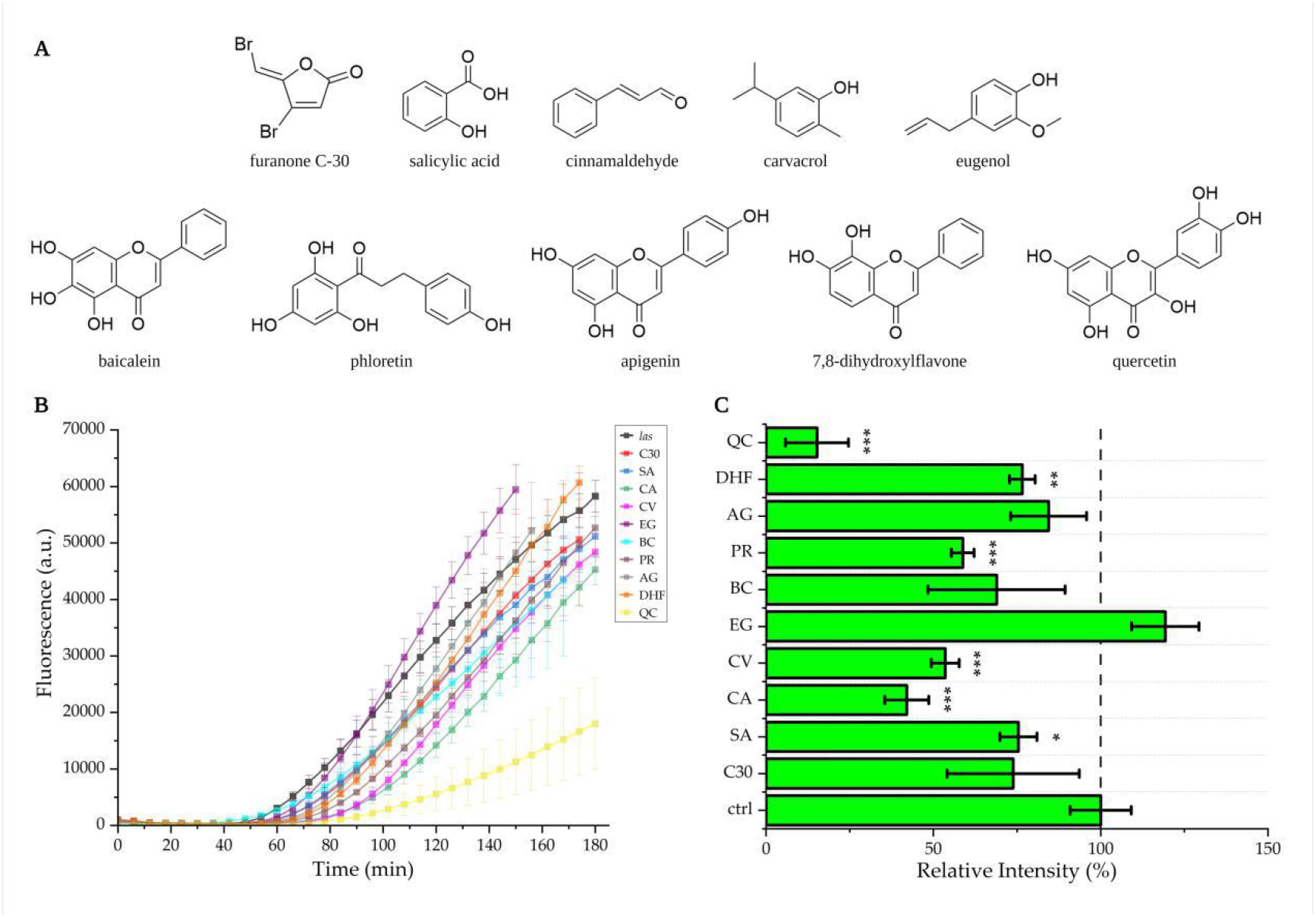
Rapid evaluation of quorum sensing inhibitors. (A) 10 quorum sensing inhibitors used in this work. (B) Fluorescence kinetics of *las* pathways treated by 10 QSIs.C30 (red) denotes furanone C-30; SA (blue) denotes salicylic acid; CA (green) denotes cinnamaldehyde; CV (magenta) denotes carvacrol; EG (purple) denotes eugenol; BC (cyan) denotes baicalein; AG (silver) denotes apigenin; PR (brown) denotes phloretin; DHF (orange) denotes 7,8-dihydroxylflavone; QC (yellow) denotes quercetin. (C) Inhibitory effects of the QSIs posed to *las* pathway. The data represent measurements taken at 2 h, with the background fluorescence subtracted. Statistical significance was determined using unpaired two-tailed Student’s *t* tests. ***, p<0.005; **, p<0.01; *, p<0.05. The experimental data are presented as mean (s.d.) from sample sizes of n=3 biological replicates drawn from a single batch of *E. coli* extract.

Because the *rhl* pathway functioned unexpectedly weak with highly fluctuating outcome, we chose the more robust *las* pathway for proof-of-concept. Candidate QSIs were added to the cell-free *las* pathway at recommended sub-inhibitory concentrations for evaluation [9,13-15,44-47]: 10 μM furanone C-30; 1.5 mM salicylic acid; 1.5 mM cinnamaldehyde; 200 μM eugenol; 1 mM carvacrol; 100 μM baicalein; 100 μM quercetin; 100 μM apigenin; 100 μM phloretin; and 100 μM 7,8-dihydroxylflavone. In a reconstituted *las* pathway, the QS components (LasI, LasR and 3-oxo-C12-HSL) were continuously synthesized, accumulated and saturated, while the QSIs were only added at the beginning, rendering it difficult to quantify the inhibitory effects of QSIs by endpoint measurements (Figure S5). Therefore, considering the exceptionally rapid reaction kinetics of cell-free expression, we focused on early-stage fluorescence monitoring to evaluate the inhibitory efficacy of candidate molecules. Notably, considerable discrepancy in fluorescence expression was found during 3-hour kinetics detection, enabling rapid, real-time report and analysis (Figure 2B). 9/10 of tested molecules displayed mild to high inhibition to cell-free *las* pathway at 2 hour (Figure 2C), except eugenol, which surprisingly reinforced the activity of cell-free *las* pathway. Of potent molecules, quercetin displayed the highest inhibition (84.8 ± 9.4%) and markedly delayed pLas-driven GFP expression, consistent with its potential mechanisms of noncompetitive binding to LasR for allosteric modulation and transcription inhibition [14]. Also, phloretin, carvacrol and cinnamaldehyde showed significant inhibition (p<0.01). Furanone C-30 was considered as a positive control of the evaluation, and inhibited the pathway at 10 μM as expected. Yet, the efficacy of other tested flavonoid QSIs, *e*.*g*., baicalein [14,45] and 7,8-dihydroxylflavone [14], greatly differed with previous research using *E. coli* reporter strains, in which they reduced bioluminescence production by 70%∼80%. We speculate this discrepancy might stem from the the use of different pLas promoters. More specifically, flavonoid QSIs might selectively interfere with the interactions between a LuxR-type transcription factor and its cognate *cis*-regulatory components, and in-depth investigation about how flavonoid QSIs impact gene expression driven by varied pLux-type promoters is supposed to give more insights into their inhibitory mechanisms, such as identifying particular *cis*-motifs responsible for signal blocking.

### Quantification of on-target inhibition and off-target toxicity

The conventional methods tend to take bacterial growth (represented by OD_600_) as the metric to evaluate candidate QSI’s nonspecific toxicity to the cell, so as to overcome the false-positive dilemma. Notwithstanding low growth inhibition reported for most QSIs, we doubt whether measuring bacterial density is enough to infer the off-target toxicity, thinking living bacterium as a robust system tolerant of changeable surroundings, and to further eliminate the possibility of false-positive. Importantly, in contrast to living reporter bacteria with a plethora of understudied (yet possibly implicated) factors and mechanisms, cell-free quorum sensing pathways function only through *in vitro* transcription/translation, enzyme catalysis and ligand-receptor interaction, minimizing misleading targets and thus suited for accurate drug evaluation.

To more comprehensively evaluate the QSIs, we defined “on-target inhibition” as their specific interference with quorum sensing components, “off-target toxicity” as their nonspecific damage to cell-free expression machinery. To experimentally distinguish on-target inhibition from off-target toxicity, we supplemented the QSIs into cell-free reactions expressing GFP by a σ_70_-driven constitutive promoter. The 4-hour dynamic results (Figure 3A) showed that 6/10 of candidate QSIs exhibited little or mild off-target toxicity in previously tested concentrations, while salicylic acid, carvacrol, quercetin and cinnamaldehyde moderately reduced GFP expression (p<0.01, Figure 3B). The endpoint measurements substantiated that cinnamaldehyde at 1.5 mM damaged cell-free expression machinery (Figure S6). Interestingly, salicylic acid caused similar extent of GFP expression reduction in the assays quantifying on-target inhibition (24.6 ± 5.5%) and off-target toxicity (25.4 ± 2.0%), making us doubt whether this phytohormone specifically inhibits LasR as previous *in silico* prediction suggested [13]. Baicalein, 7,8-dihydroxylflavone and apigenin showed negligible off-target toxicity as well as a certain level of on-target inhibition, analogous to furanone C-30, and were thus considered as potential targeted anti-QS agents. Moreover, phloretin attenuated cell-free *las* pathway at 41.2 ± 3.4% and constitutive expression at 16.5 ± 7.6%, being evaluated as a potent QSI candidate.

**Figure 3:**
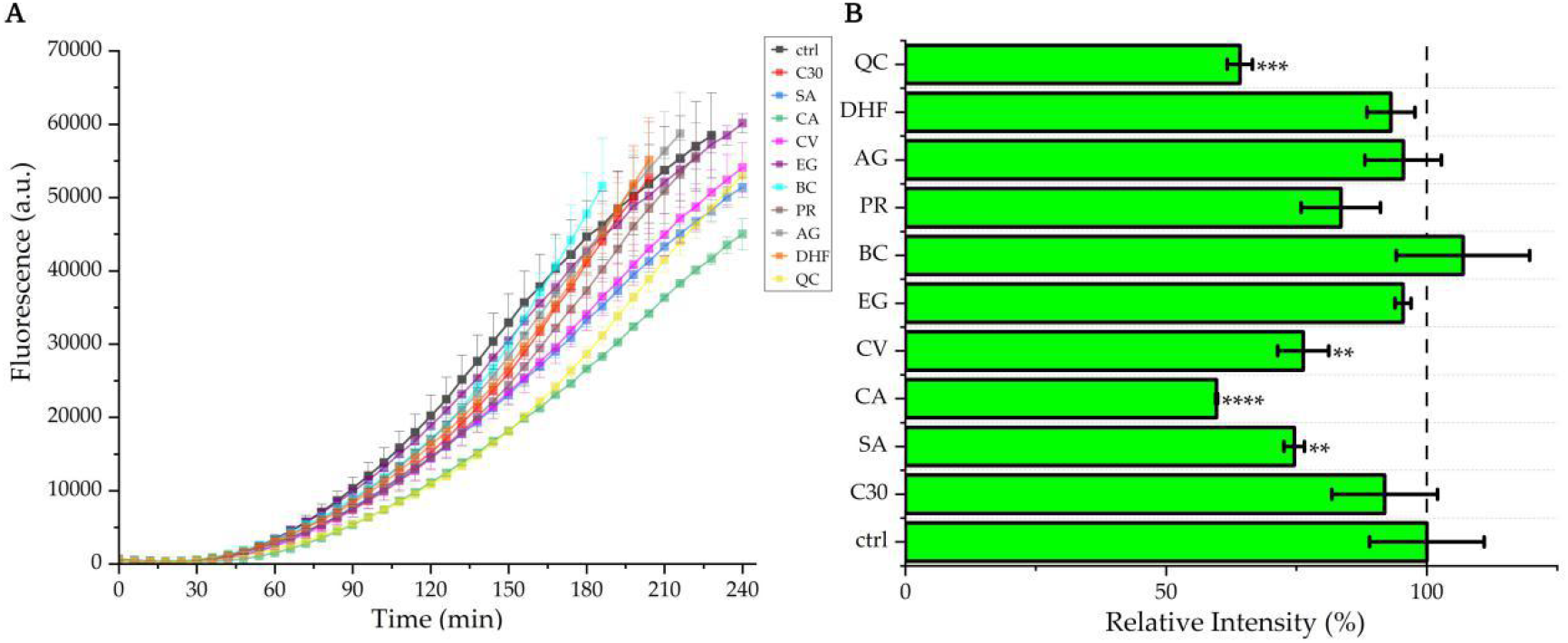
Off-target toxicity of select QSIs. (A) 4-hour fluorescence kinetics of the cell-free reactions constitutively expressing GFP (5 nM pDY5) treated by ten QSIs. Off-target toxicity of ten compounds to the cell-free expression machinery was quantified by inhibition of constitutive GFP expression. (B) Cinnamaldehyde, quercetin, carvacrol and salicylic acid posed moderate interference to cell-free expression machinery. The data represent measurements taken at 3 hour, with the background fluorescence subtracted. Statistical significance was determined using unpaired one-tailed Student’s *t* tests. ****, p<0.0001; ***, p<0.001; **, p<0.01. The experimental data are presented as mean (s.d.) from sample sizes of n=3 biological replicates drawn from a single batch of *E. coli* extract.

Given its prominent inhibitory efficacy towards the cell-free *las* pathway, we further investigated the dose response of quercetin. Quercetin in a concentration gradient (0, 10 μM, 20 μM, 50 μM, 100 μM, 200 μM) was supplemented to cell-free reactions either reconstituting *las* pathway or constitutively expressing GFP. The results showed that quercetin displayed dose-dependent inhibition towards *las* pathway (Figure 4A), and acceptable off-target toxicity (20%∼30%) within 100 μM (Figure 4B). Importantly, quercetin significantly delayed the QS response in the kinetics assays (Figure 4A), even at 10 μM. Upon the rapid evaluations about on-target inhibition and off-target toxicity, quercetin was demonstrated as a promising QSI candidate that specifically targets *las* pathway of *P. aeruginosa*.

**Figure 4:**
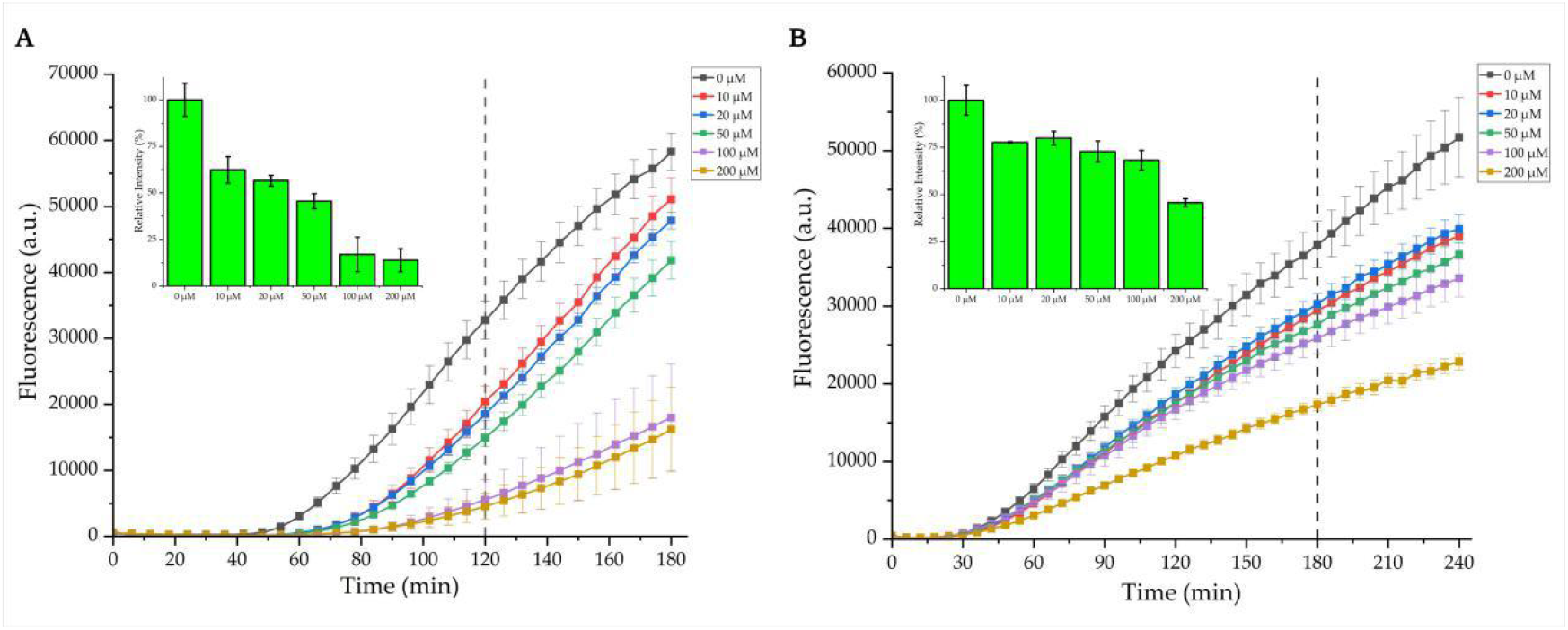
Amongst the candidate QSIs, quercertin exhibited (A) dose-dependent on-target inhibition, and (B) acceptable off-target toxicity. The experimental data are presented as mean (s.d.) from sample sizes of n=3 biological replicates.

## CONCLUSIONS

With an increasing number of microbial quorum sensing systems being reported, synthetic biologists are keen to excavate, optimize and standardize QS components, *e*.*g*., enzymes responsible for AI synthesis, receptors that capture AIs sensitively, and AI-responsive promoters for expressing gene of interest with broad dynamic range, and to design artificial quorum sensing circuitry with high orthogonality. Notably, the synthetic biology community has reprogrammed quorum sensing as a toolkit to rationally regulate cellular group activities, facilitating myriad applications including population density control [48,49], biological pattern formation [50,51] and edge detection [52], biosensing [53], biocomputing [54,55], growth-coupled metabolic engineering [56], social interaction programming [57], and solid tumor therapy mediated by quorum oscillators [58,59].

Despite above-mentioned remarkable advances, the biological significance of quorum sensing systems should not be overlooked. In this study, focusing on pharmaceutical potential of quorum sensing, we *in vitro* reconstituted two quorum sensing pathways of an important human pathogen using cell-free expression techniques, and quantitatively investigated the efficacy of previously-reported quorum sensing inhibitors on the *las* pathway within hours. Of ten plant-derived molecules, most flavonoid candidates specifically inhibited the cell-free *las* pathway, consistent with previous reports, especially quercetin, which was later characterized to display potent, dose-dependent inhibition and acceptable off-target toxicity at low concentrations. Yet, salicylic acid did not show particular inhibition and eugenol even accelerated the kinetics of the reconstituted *las* pathway, questioning whether they indeed target LasR for inhibition as reported before, and suggesting other cryptic mechanisms responsible for their observed inhibitory efficacy *in vivo*. In addition, we propose that the compounds that display considerable off-target toxicity, such as cinnamaldehyde and carvacrol, might not be evaluated properly, revealing a pitfall of the methodology and asking further optimization of cell-free expression robustness.

A wide array of QSIs are hypothesized to target specific QS components. Yet, for most of them, the inhibitory mechanisms remain at the stage of *in silico* prediction and lack reasonable experimental proof, thus susceptible to false-positive. We envision that our cell-free QS pathways could be utilized for exploring the inhibitory mechanisms of QSIs, *e*.*g*., whether the QSIs impair the catalytic activity of autoinducer synthases. For *in vivo* research, despite the reduction of AI synthesis observed, the enzymatic reactions inside bacteria are in fact complicated by unelucidated, higher-order regulatory networks. Combining cell-free enzymatic reactions with LC-MS, one might readily figure out whether a QSI candidate inhibits the biosynthesis of autoinducers. We believe the *in vitro* platform proposed herein offers a route complementary to *in vivo* QSI evaluation, and the data gleaned from cell-free pathways could rationally instruct *in vivo* study. Hopefully, the versatility of cell-free drug-evaluation platform can be further explored by rewiring other druggable bacterial signal transduction pathways.

Cell-free expression platform holds great promise for advancing biology and biotechnology, owing to its exceptional flexibility and modularity. Firstly, bypassing cellular barriers, it allows scientists to manipulate genetic materials only by liquid handling, which is highly compatible with automatic experimentation platforms, such as acoustic liquid handling robotics [28] or microfluidic chips [27], and simultaneously ensures reproducibility when performing miniaturized reactions. Secondly, bypassing most cryptic endogenous factors hidden in the genome, it offers a reductionist perspective to reconstitute cellular biophysical and biochemical process *in vitro* and to potentiate methodological innovations. Inspired by these merits, our prospective work aims to facilitate reliable, rapid cell-free drug screening in high throughput, contrasting conventional *in vivo* drug-screening methods, which are usually precluded by limited throughput, time-consuming experimental cycle, and occasionally identify off-target candidates. Recent developments in deep learning have been reshaping the fields of drug discovery. Of note, Stokes *et al*. [60] screened 2,335 diverse molecules in an *Escherichia coli* growth inhibition assay, and utilized generated datasets to train a deep neural network model for identifying novel antibiotics from >107 million records in chemical databases. Intriguingly, will datasets generated from rationally designed cell-free pathways, other than from living cells, be able to train deep-learning algorithms that enable accurate predictions with customizable drug targets? In the face of formidable antimicrobial resistance crisis, we anticipate our cell-free drug-evaluation platform is capable to bridge *in silico* and *in vivo* drug-screening methods, and accelerate the upgrading of antimicrobial arsenal.

## METHODS

### Plasmids construction and purification

DNA used in the study was synthesized by Genscript and IDT, and was cloned into plasmid backbone pSB1A3 (iGEM distribution kit) via gibson assembly, generating pDY5, pDY7, pDY9, pDY10, pDY11, pDY13 (see Supporting Information for detailed plasmid architectures). pJL1 was a gift from Prof. M. C. Jewett (Northwestern University). *Escherichia coli* DH10β was used for plasmids cloning and amplification. All plasmids were purified by Axygen® AxyPrep Plasmid Kit (New York, USA) for cell-free expression.

### *Escherichia coli* Lysates Preparation

*E. coli* Star™ BL21(DE3) was purchased from Tsingke Biotechnology (Wuhan, China) for lysates preparation. We suspect this strain with lower RNase activity is a better starting chassis, since enhanced mRNA stability is a key factor to enable efficient cell-free protein synthesis using bacterial transcription system [37]. The protocol is mainly adapted from Jewett and Swartz [36], and modified according to literature [28,32,37-40] to activate endogenous components *in vitro. E. coli* cells were grown in 1L 2×YTP medium (10 g/L yeast extract, 16 g/L tryptone, 5 g/L sodium chloride, 7 g/L potassium phosphate dibasic, 3 g/L potassium phosphate monobasic) in a 2.5-L shake flask, at 37 °Cand 250 rpm. When OD_600_ reached 0.6, IPTG (0.1 mM) was added (using T7-driven GFP expression as quality control) and temperature was set to 30 °C. When OD_600_ reached 3.0, cells were harvested by centrifugation at 5000 g and 4 °Cfor 15 min. The cell pellets were washed by 25 mL S12 buffer (10 mM Tris, 14 mM magnesium acetate, 60 mM potassium glutamate, 2 mM DTT, brought to pH 8.2 by acetic acid) for thrice and centrifuged at 5000 g and 4 °Cfor twice. After the final centrifugation at 7000 g and 4 °Cfor 10 min, cell pellets were re-suspended in 1 mL S12 buffer/g pellets and lysed by sonication (Vibra-Cell™, Sonics). The sonication was performed at 20 kHz frequency and 50% amplitude by 10s ON/ 40s OFF pulses for delivering ∼500 J/mL cell suspensions. The lysates were then centrifuged at 12000 g and 4 °Cfor 15 min, and the supernatant was removed into 1.5-mL tubes by pipetting. The tubes containing crude *E. coli* lysates were incubated at 220 rpm for 80 min at 37 °Cor 4 hours at 30 °C. After the “ribosomal run-off” reaction, the turbid lysates were re-centrifuged at 12000 g and 4 °Cfor 15 min, and the supernatant was removed, aliquoted, flash-frozen in liquid nitrogen and stored at −80 °Cuntil use.

### Cell-Free Expression Reactions

A typical cell-free expression reaction in this study consisted of 25% (v/v) 4×premix solution (4.8 mM ATP; 3.4 mM each of GTP, UTP, and CTP; 136.0 μg/mL folinic acid; 680 μg/mL *E. coli* tRNA; 520 mM potassium glutamate; 40 mM ammonium glutamate; 48 mM magnesium glutamate; 6 mM each of 20 amino acids; 1.32 mM nicotinamide adenine dinucleotide; 1.08 mM coenzyme A; 6 mM spermidine; 4 mM putrescine; 16 mM sodium oxalate; 132 mM phosphoenolpyruvate), and 30% (v/v) *E. coli* lysates, plasmids and mother solutions of select QSIs (Supplementary Methods) to the desired concentration, and ddH_2_O to make up the volume.

For fluorescence measurement, reactions were prepared in centrifuge tubes (one-pot reactions at 10-μL scale; two-step reactions at 12.5-μL scale) and transferred to a black, 96-well microplate (Greiner-Bio One, 675077), each condition in triplicate. Plates were sealed by MonAmp™ Ultraclear Peelable Films (Monad, MQ65001) and GFP fluorescence (emission/excitation: 485/525 nm) was detected by microplate readers (Infinite M Plex, Tecan; SpectraMax i3x, Molecular Devices). For fluorescence kinetics, the reactions were monitored every 6 min for 3 or 4 hours at 30 °Cusing Infinite M Plex. For endpoint experiments, the reactions were incubated at 30 °Cfor 16 hours, and 4 μL reaction solution was transferred to 96-well plate, diluted to 100 μL by water, and measured by SpectraMax i3x. For LC-MS analysis, we prepared 100-μL reactions in 1.5-mL eppendorf tubes, and incubated them for 16 hours at 30 °Cand 220 rpm.

### LC-MS Analysis

To extract synthesized AHL from reactions, we pipetted 100 μL ethyl acetate (acidified by 0.1% formic acid) into each overnight reaction. The reactions were gently vortexed for 5 min, and centrifuged at 3000 g for 3 min, after which the top-layer organic phase was carefully removed. The process was repeated for three times, and the collected samples were freeze-dried and re-suspended by 100 μL methanol for later analysis.

LC-MS analysis was conducted with Dionex UltiMate 3000 UHPLC system (Thermo Scientific) and Ultra High Resolution Linear Ion Trap Orbitrap Mass Spectrometer (Thermo Scientific). A Hypersil GOLD C18 aQ (150 mm × 2.1 mm, Thermo Scientific) column was used for LC, with water and acetonitrile as mobile phase (acidified by 0.1% formic acid). Standard samples of 3-oxo-C12-HSL and C4-HSL were purchased from Sigma-Aldrich. The cell-free reaction samples were preprocessed using HPLC (Supplementary Methods). The detailed LC-MS data were documented in Supporting Information (Figure S3, S4).

### Data Analysis and Statistics

All data were analyzed and plotted with Origin, Version 2021b (OriginLab Corporation, Northampton, MA, USA). Standard curves that convert relative fluorescence unit (RFU) to GFP concentration were illustrated using SpectraMax i3x (Figure S1), and the endpoint values measured by the microplate reader were normalized using the standard curve.

The figures were illustrated and assembled using BioRender.com.

## ACKNOWLEDGEMENTS

This work was supported by the National Key R&D Program of China [grant number 2018YFA0900400] and the National Natural Science Foundation of China [grant number 31971341].

## Notes

### Competing Interest Statement

The authors have declared no competing interest.

