## Supporting Information for "In vitro reconstituted quorum sensing pathways enable rapid evaluation of quorum sensing inhibitors" (from another version of this preprint: version 1, 2021-10-29)

Dingchen Yu[1]*

[1]College of Life Sciences, Wuhan University, Wuhan 430072, PR China

### Supporting Information

#### Table of Contents


| **Supplementary Material and Methods** | **Page** |
|---|---|
| DNA part sequences | 2 |
| Plasmid maps | 4 |
| Preparation of quorum sensing inhibitors | 10 |
| Preprocessing of cell-free reaction samples with HPLC | 11 |
| **Supplementary Figures** | |
| Figure S1: Standard curve quantifying GFP expression generated with SpectraMax i3x | 12 |
| Figure S2: Cell-free systems based on previously- and currently-used protocols vary in TXTL performance | 13 |
| Figure S3: LC-MS analysis of cell-free *las* pathway | 14 |
| Figure S4: LC-MS analysis of cell-free *rhl* pathway | 15 |
| Figure S5: Endpoint measurements of cell-free *las* pathways treated by QSIs | 16 |
| Figure S6: Endpoint measurements of cell-free reactions constitutively expressing GFP treated by QSIs | 17 |
| Figure S7: Endpoint measurements of cell-free *las* pathways treated by quercetin with varied concentrations | 18 |
| Figure S8: Endpoint measurements of cell-free reactions constitutively expressing GFP treated by quercetin with varied concentrations | 19 |
| **References** | 20 |

### Supplementary Material and Methods

#### DNA part sequences

Table of part sequences used in pDY5, pDY7, pDY9, pDY10, pDY11 and pDY13. Names beginning with BBa_ refer to Part Numbers that were obtained from the BioBricks Parts Registry (http://parts.igem.org/Main_Page).

| Name | Part type | Sequence |
|---|---|---|
| BBa_J23100 | Promoter | ttgacggctagctcagtcctaggtacagtgctagc |
| pLas<br>BBa_K649000 | Promoter | ttcgagcctagcaagggtccgggttcaccgaaatctatctcatttgctagttataaaatt<br>atgaaatttgcgtaaattcttca |
| pRhl<br>BBa_K1949060 | Promoter | tcctgtgaaatctggcagttaccgttagctttcgaattggctataaagtgttc |
| BBa_B0030 | RBS | attaaagaggagaaa |
| BBa_B0034 | RBS | aaagaggagaaa |
| BBa_B1002 | Terminator | cgcaaaaaaccccgcttcggcggggttttttcgc |
| LasR<br>BBa_C0179 | CDS | atggccttggttgacggttttcttgagctggaacgctcaagtggaaaattggagtggagc<br>gccatcctccagaagatggcgagcgaccttggattctcgaagatcctgttcggcctgttg<br>cctaaggacagccaggactacgagaacgccttcatcgtcggcaactacccggccgcctgg<br>cgcgagcattacgaccgggctggctacgcgcgggtcgacccgacggtcagtcactgtacc<br>cagagcgtactgccgattttctgggaaccgtccatctaccagacgcgaaagcagcacgag<br>ttcttcgaggaagcctcggccgccggcctggtgtatgggctgaccatgccgctgcatggt<br>gctcgcggcgaactcggcgcgctgagcctcagcgtggaagcggaaaaccgggccgaggcc<br>aaccgtttcatagagtcggtcctgccgaccctgtggatgctcaaggactacgcactgcaa<br>agcggtgccggactggccttcgaacatccggtcagcaaaccggtggttctgaccagccgg<br>gagaaggaagtgttgcagtggtgcgccatcggcaagaccagttgggagatatcggttatc<br>tgcaactgctcggaagccaatgtgaacttccatatgggaaatattcggcggaagttcggt<br>gtgacctcccgccgcgtagcggccattatggccgttaatttgggtcttattactctctaa<br>taa |
| RhlR<br>BBa_C0171 | CDS | atgaggaatgacggaggcttttttgctgtggtgggacggtttgcgtagcgagatgcagccg<br>atccacgacagccagggcgtgttcgccgtcctggaaaaggaagtgcggcgcctgggcttc<br>gattactacgcctatggcgtgcgccacacgattcccttcacccggccgaagaccgaggtc<br>catggcacctatcccaaggcctggctggagcgataccagatgcagaactacggggccgtg<br>gatccggcgatcctcaacggcctgcgctcctcggaaatggtggtctggagcgacagcctg<br>ttcgaccagagccggatgctctggaacgaggctcgcgattggggcctctgtgtcggcgcg<br>accttgccgatccgcgcgccgaacaatttgctcagcgtgctttccgtggcgcgcgaccag<br>cagaacatctccagcttcgagcgcgaggaaatccgcctgcggctgcgttgcatgatcgag<br>ttgctgacccagaagctgaccgacctggagcatccgatgctgatgtccaacccggtctgc<br>ctgagccatcgcgaacgcgagatcctgcaatggaccgccgacggcaagagttccggggaa<br>atcgccatcatcctgagcatctccgagagcacggtgaacttccaccacaagaacatccag<br>aagaagttcgacgcgccgaacaagacgctggctgccgcctacgccgcggcgctgggtctc<br>atctaataa |
| LasI<br>BBa_C0178 | CDS | atgatcgttcagatcggtcgtcgtgaagagttcgacaaaaaactgctgggtgaaatgcac<br>aaactgcgtgctcaggttttcaaagaacgtaaaggttgggacgtttccgttatcgacgaa |

| | | |
|---|---|---|
| | | atggaaatcgacggttacgacgctctgtccccgtactacatgctgatccaggaagacacc<br>ccggaagctcaggttttcggttgctggcgtatcttcgacaccaccggtccgtacatgctg<br>aaaaacaccttcccggaactgctgcacggtaaagaagctccgtgctccccgcacatctgg<br>gaactgtcccgtttcgctatcaactccggtcagaaaggttccctgggtttctccgactgc<br>accctggaagctatgcgtgctctggctcgttactccttgcagaacgacatccagaccctg<br>gttaccgttaccaccgttggtgttgaaaaaatgatgatccgtgctggtctggacgtttcc<br>cgtttcggtccgcacctgaaaatcggtatcgaacgtgctgttgctctgcgtatcgaactg<br>aacgctaaaacccagatcgctctgtacggtggtgttctggttgaacagcgtctggctgtt<br>tcctaataa |
| RhlI<br>BBa_C0170 | CDS | atgatcgaactgctgtccgaatccctggaaggtctgtccgctgctatgatcgctgaactg<br>ggtcgttaccgtcaccaggttttcatcgaaaaactgggttgggacgttgtttccacctcc<br>cgtgttcgtgaccaggagttcgaccagttcgaccacccgcagacccgttacatcgttgct<br>atgtcccgtcagggtatctgcggttgcgctcgtctgctgccgaccaccgacgcttacctg<br>ctgaaagacgttttcgcttacctgtgctccgaaaccccgccgtccgacccgtccgtttgg<br>gaactgtcccgttacgctgcttccgctgctgacgacccgcagctggctatgaaaatcttc<br>tggtcctccctccagtgcgcttggtacctgggtgcttcctccgttgttgctgttaccacc<br>accgctatggaacgttacttcgttcgtaacggtgttatcctccagcgtctgggtccgccg<br>cagaaagttaaaggtgaaaccctggttgctatctccttcccggcttaccaggaacgtggt<br>ctggaaatgctgctgcgttaccacccggaatggctccagggtgttccgctgtccatggct<br>gtttaataa |
| GFPmut3b<br>BBa_E0040 | CDS | atgcgtaaaggagaagaacttttcactggagttgtcccaattcttgttgaattagatggt<br>gatgttaatgggcacaaattttctgtcagtggagagggtgaaggtgatgcaacatacgga<br>aaacttacccttaaatttatttgcactactggaaaactacctgttccatggccaacactt<br>gtcactactttcggttatggtgttcaatgctttgcgagatacccagatcatatgaaacag<br>catgactttttcaagagtgccatgcccgaaggttatgtacaggaaagaactatatttttc<br>aaagatgacgggaactacaagacacgtgctgaagtcaagtttgaaggtgatacccttgtt<br>aatagaatcgagttaaaaggtattgattttaaagaagatggaaacattcttggacacaaa<br>ttggaatacaactataactcacacaatgtatacatcatggcagacaaacaaaagaatgga<br>atcaaagttaacttcaaaattagacacaacattgaagatggaagcgttcaactagcagac<br>cattatcaacaaaatactccaattggcgatggccctgtccttttaccagacaaccattac<br>ctgtccacacaatctgccctttcgaaagatcccaacgaaaagagagaccacatggtcctt<br>cttgagtttgtaacagctgctgggattacacatggcatggatgaactatacaaataataa |

**Plasmid maps**

Plasmid map for pDY5

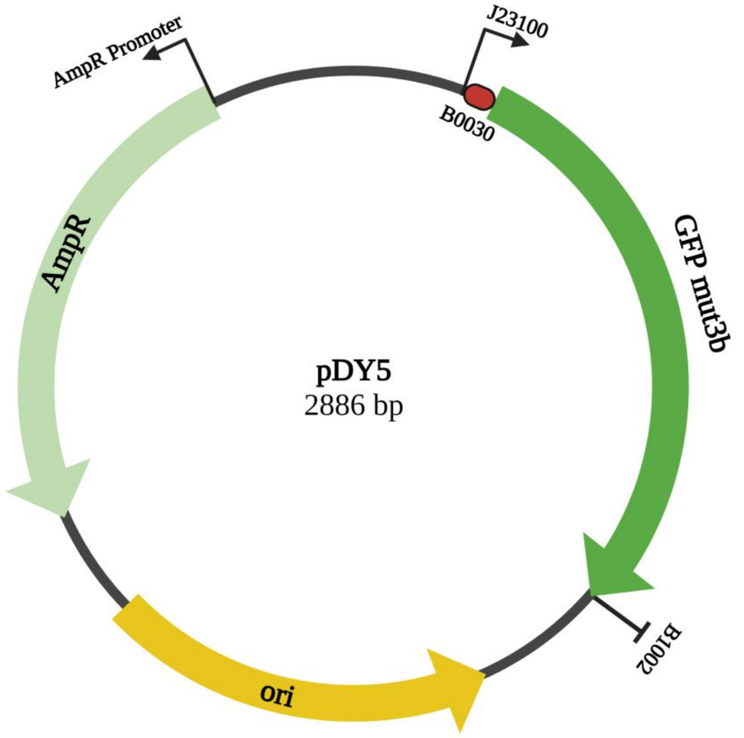

Plasmid map for pDY7

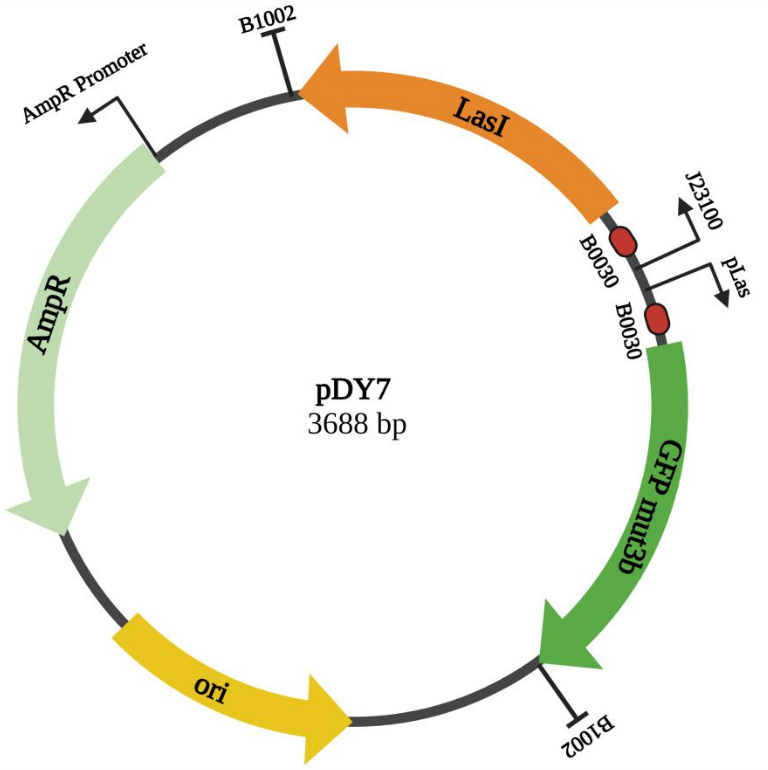

Plasmid map for pDY9

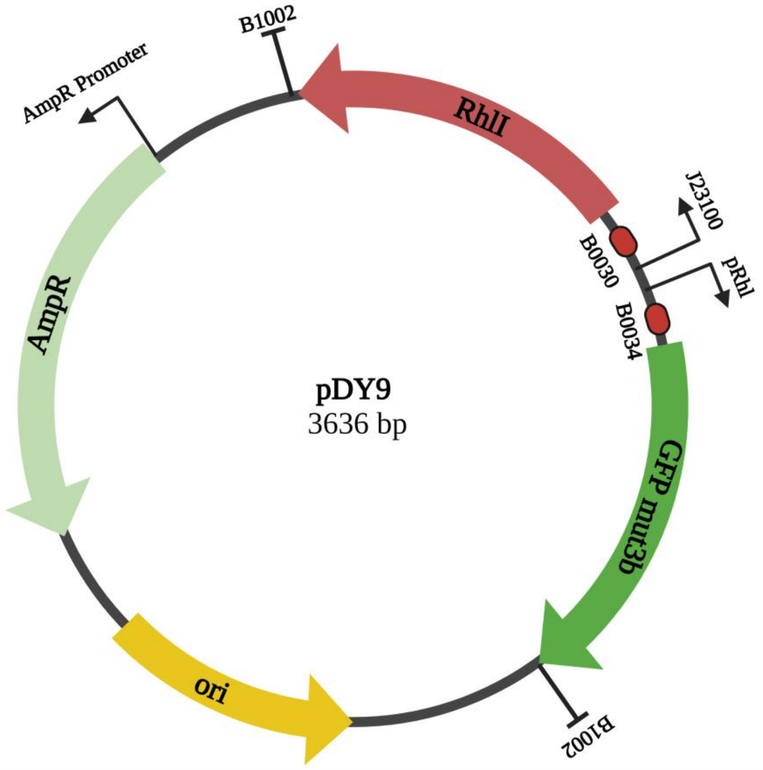

Plasmid map for pDY10

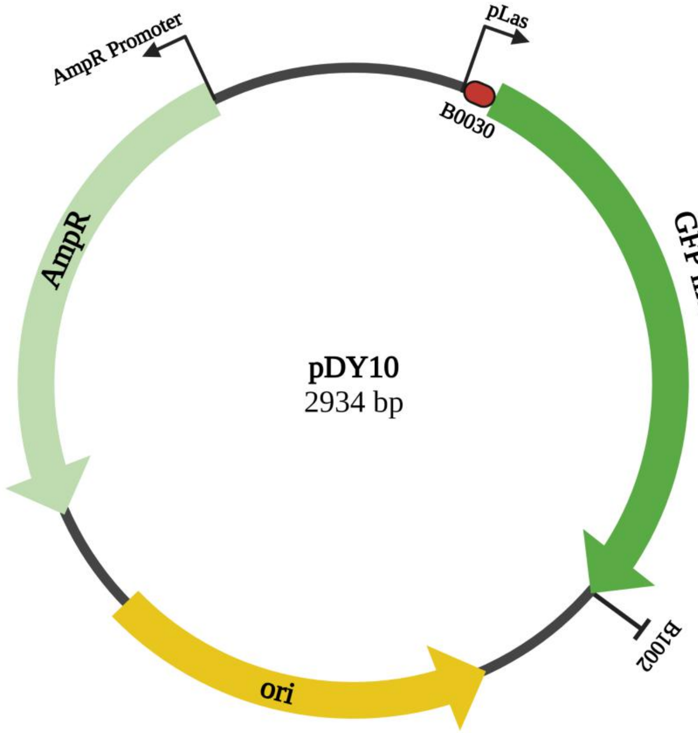

Plasmid map for pDY11

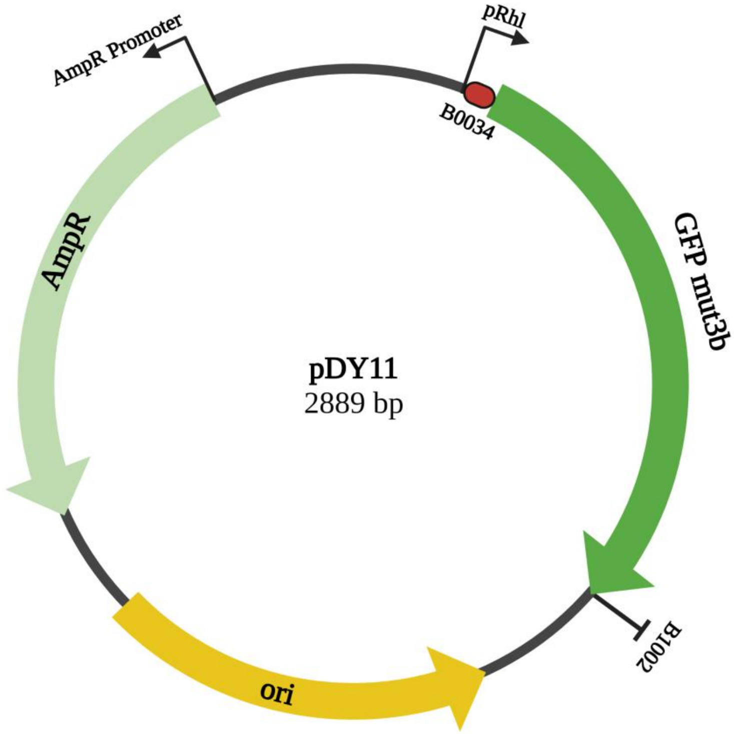

Plasmid map for pDY13

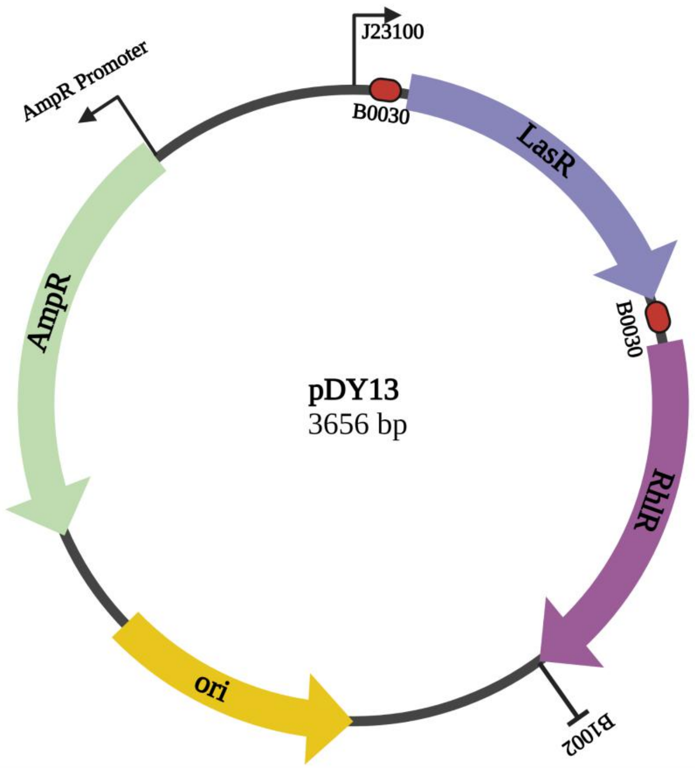

**Preparation of quorum sensing inhibitors**

Salicylic acid was purchased from Yuanye (Shanghai, China). Cinnamaldehyde was purchased from Macklin (Shanghai, China). Furanone C-30 was purchased from Sigma-Aldrich (St.Louis, US). Eugenol, carvacrol, apigenin, baicalein, phloretin, quercetin and 7,8-dihydroxylflavone were purchased from Aladdin (Shanghai, China). We used distilled Milli-Q water or akalescent S12 buffer (pH≈8.2) as solvent, because: (i) we found that bacterial native expression machinery displays low tolerance of commonly-used solvents, such as methanol, ethanol and DMSO; (ii) the optimal solvent to each compound varies, confusing the design of negative control groups and ensuing analysis; (iii) cell-free expression system is aqueous, thus, introducing compounds dissolved by organic solvents into CFE system might cause solutes precipitation without knowing. The following table briefly documents how we prepared ten chemically diverse quorum sensing inhibitors for evaluation in cell-free *las* pathway.

| Quorum sensing inhibitors | Methods |
|---|---|
| Salicylic acid | 10 mM mother solution: Add 1.4 mg salicylic acid powder into 1 mL water; Boil till the powder dissolves; Store at room temperature. |
| Cinnamaldehyde | 10 mM mother solution: Pipette 1.26 μL liquid cinnamaldehyde into 1 mL water; Store at room temperature and avoid light; Vortex before use. |
| Furanone C-30 | 250 μM mother solution: Add 1.3 mg furanone C-30 powder into 20 mL S12 buffer; Sonicate for 20 min to dissolve the powder; Store at 4℃ and avoid light. |
| Eugenol | 2 mM mother solution: Pipette 1.54 μL liquid eugenol into 5 mL water; Store at 4℃ and avoid light; Vortex before use. |
| Carvacrol | 20 mM mother solution: Pipette 3.4 μL liquid carvacrol into 1 mL water; Store at 4℃ and avoid light; Vortex before use. |
| Baicalein | 2 mM mother solution: Add 1.1 mg baicalein powder into 2 mL S12 buffer; Store at 4℃ and avoid light; Vortex before use. |
| Quercetin | 2 mM mother solution: Add 1.2 mg quecertin powder into 2 mL S12 buffer; Store at 4℃ and avoid light; Vortex before use. |
| Apigenin | 2 mM mother solution: Add 1.1 mg apigenin powder into 2 mL S12 buffer; Store at 4℃ and avoid light; Vortex before use. |
| Phloretin | 2 mM mother solution: Add 1.1 mg phloretin powder into 2 mL S12 buffer; Store at 4℃ and avoid light; Vortex before use. |
| 7,8-dihydroxylflavone | 2 mM mother solution: Add 1.3 mg 7,8-dihydroxylflavone powder into 2 mL S12 buffer; Store at 4℃ and avoid light; Vortex before use. |

**Preprocessing of cell-free reaction samples with HPLC**

There exist numerous endogenous metabolites in the *E. coli* lysates-based cell-free reactions, and unprocessed CFE samples presumably lower the accuracy of LC-MS for detecting the trace amount of AHLs within. Thus, we used Dionex UltiMate 3000 UHPLC system (Thermo Scientific) for preprocessing of cell-free *las* and *rhl* pathways, with a Hypersil GOLD C18 column (250 × 4.6 mm, Thermo Scientific). The fluids were collected to 2-mL Eppendorf tubes, with the retention time matched to standard samples of 3-oxo-C12-HSL (26.4 min-17.4 min) and C4-HSL (6.2 min-7.2 min), then freeze-dried and re-dissolved by methanol or acetonitrile for LC-MS analysis.

The HPLC protocol to roughly isolating 3-oxo-C12-HSL and C4-HSL from *E. coli* metabolome was documented as follows:

| Time | Mobile Phase | Flow Velocity |
|---|---|---|
| -5~0 min | 90% water; 10% acetonitrile | 1.0 mL/min |
| 0~10 min | 90% water; 10% acetonitrile -> 75% water; 25% acetonitrile | 1.0 mL/min |
| 10~35 min | 75% water; 25% acetonitrile -> 100% acetonitrile | 1.0 mL/min |
| 35-40 min | 100% acetonitrile | 1.0 mL/min |

### Supplementary Figures

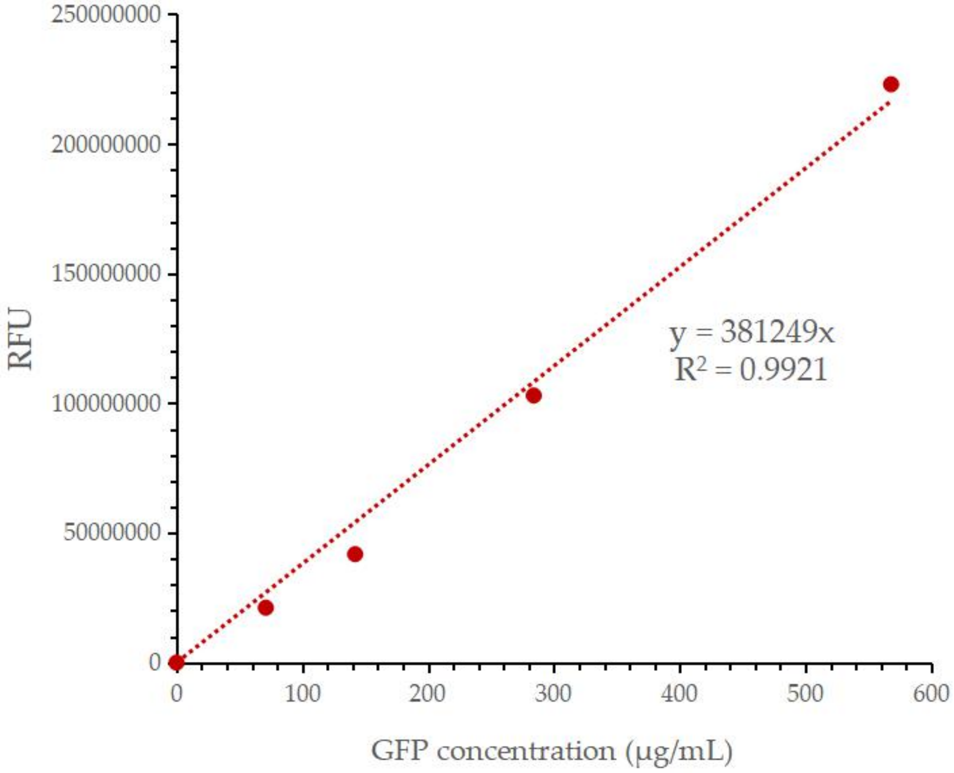


Figure S1: Standard curve quantifying GFP expression generated with SpectraMax i3x. The curve converts RFU measured by the microplate reader to GFP concentration, when implementing endpoint measurements.

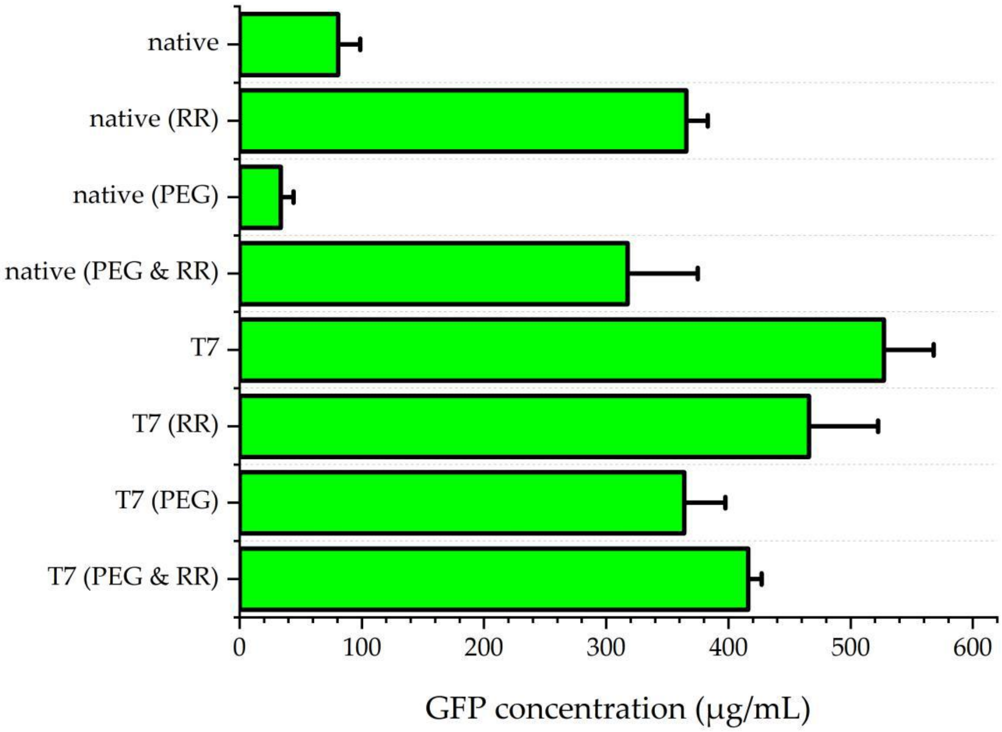


Figure S2: Cell-free systems based on previously- and currently-used protocols vary in TXTL performance. PEG denotes addition of 2.0% (w/v) PEG 8000; RR denotes the "Ribosomal Run-off" reaction. According to literature, we suspected post-lysis processing and molecular crowding agents would most likely alter the performance of our system. Artificial macromolecular crowding agents, like PEG 8000, are believed to be a vital factor to enable interactions of bacterial TXTL components *in vitro*, considering the 20-fold diluted cellular components [15]. Post-lysis incubation is also reported as necessary, since the process is believed to digest most of endogenous transcripts, sparing the ribosomes and eliminating the competitor mRNA to enable more efficient *in vitro* expression. We used the plasmids, pJL1 (5 nM) and pDY5 (5 nM), to evaluate T7 and bacterial native expression system, respectively. The T7 system maintained robust expression under different tested conditions, yielding GFP at >500 μg/mL without additional processing. Yet, for bacterial native expression machinery, appropriate ribosomal run-off reaction (80 min at 37℃ or 4 hours at 30℃) was vital. Though PEG 8000 was reported useful to enhance *B. megaterium* cell-free expression [9], we observed slight reduction of GFP expression with the addition of PEG 8000 in our *E. coli*-based system, for both native expression and T7 expression. Polyamines in the premix buffer, *i.e.*, putrescine and spermidine, seemed sufficient to enact molecular crowding [11]. Ultimately, we decide that, for T7 expression, no change is made to the protocol; for native expression, the ribosomal run-off reaction is added to our currently-used protocol.

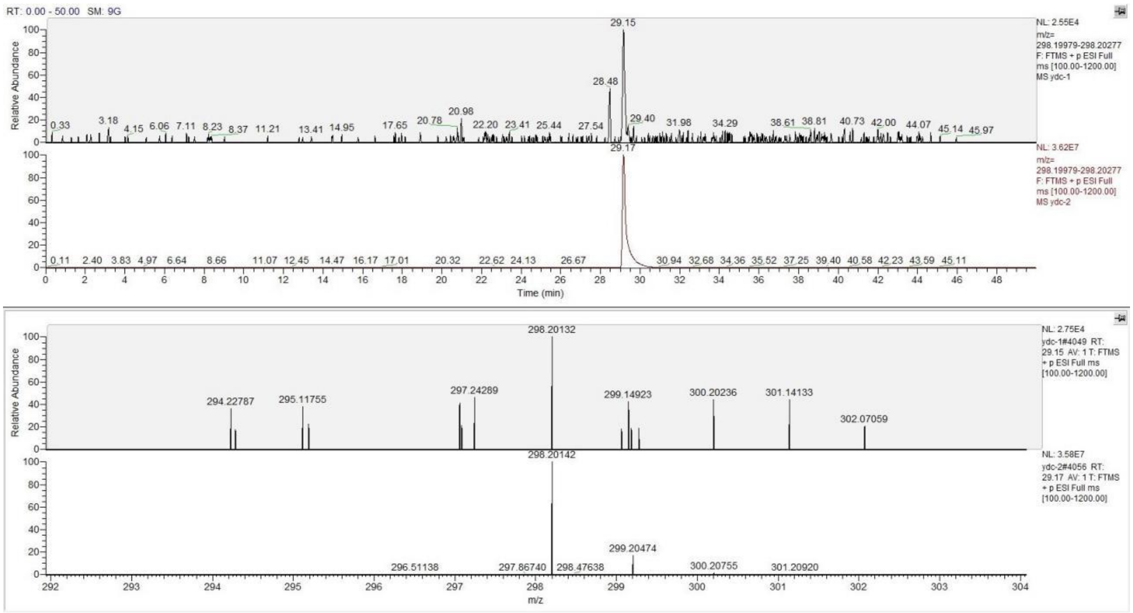


Figure S3: LC-MS analysis of cell-free *las* pathway. The LC-MS data were analyzed using Thermo Xcalibur Qual Browser. In each compartment, the top results represent cell-free reaction samples, and the bottom results represent standard samples of 3-oxo-C12-HSL (m/z = 298.201).

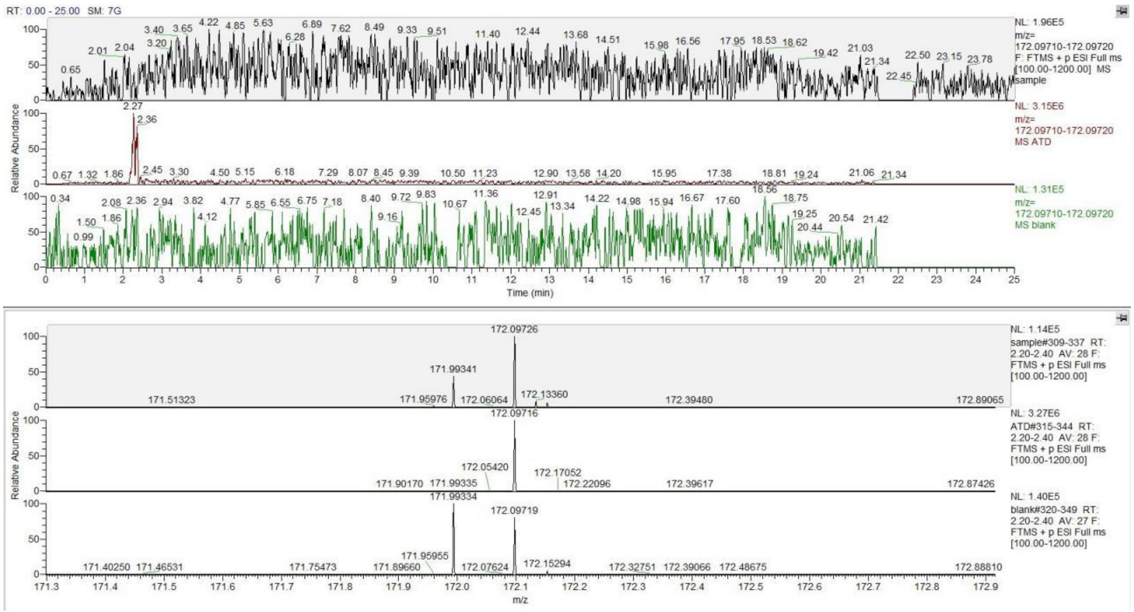


Figure S4: LC-MS analysis of cell-free *rhl* pathway. In each compartment, the top results represent cell-free reaction samples, the middle results represent standard samples of C4-HSL (m/z = 172.097), and the bottom results represent blank samples (acetonitrile).

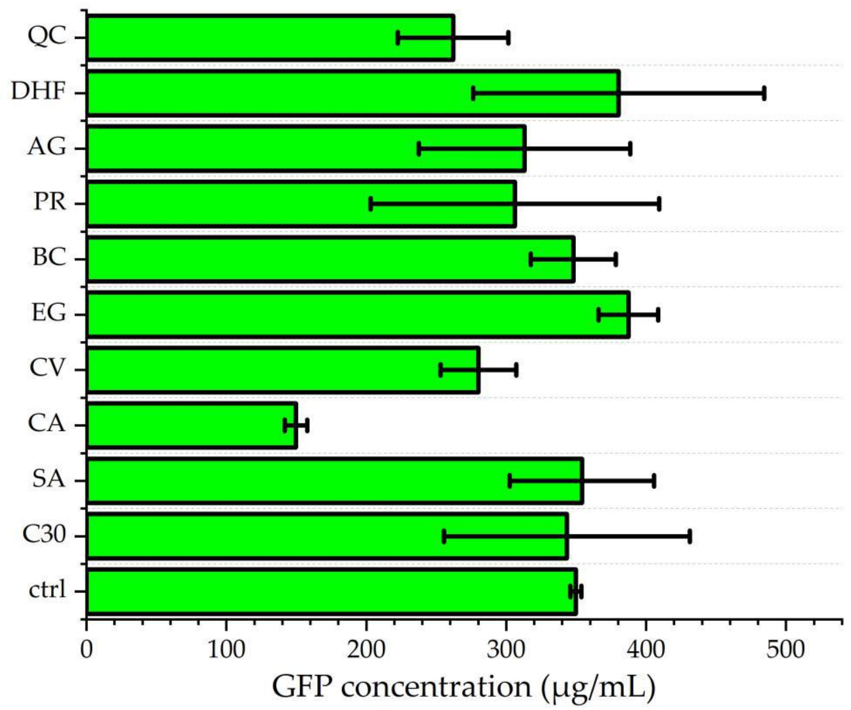


Figure S5: Endpoint measurements of cell-free *las* pathways treated by QSIs. Cinnamaldehyde, quercetin and carvacrol manifested considerable inhibition.

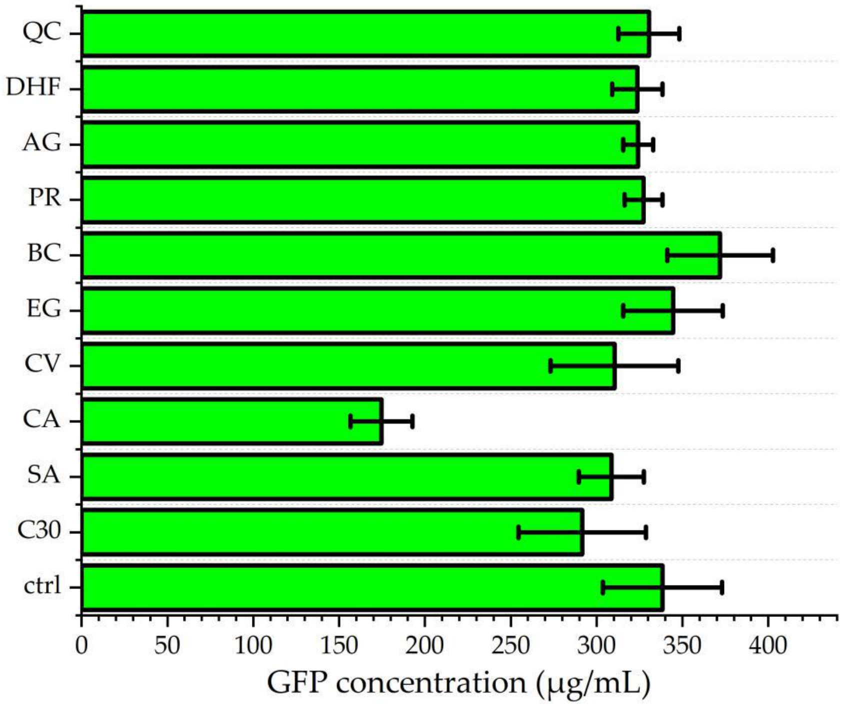


Figure S6: Endpoint measurements of cell-free reactions constitutively expressing GFP treated by QSIs. Cinnamaldehyde was the only candidate that significantly reduced constitutive GFP expression.

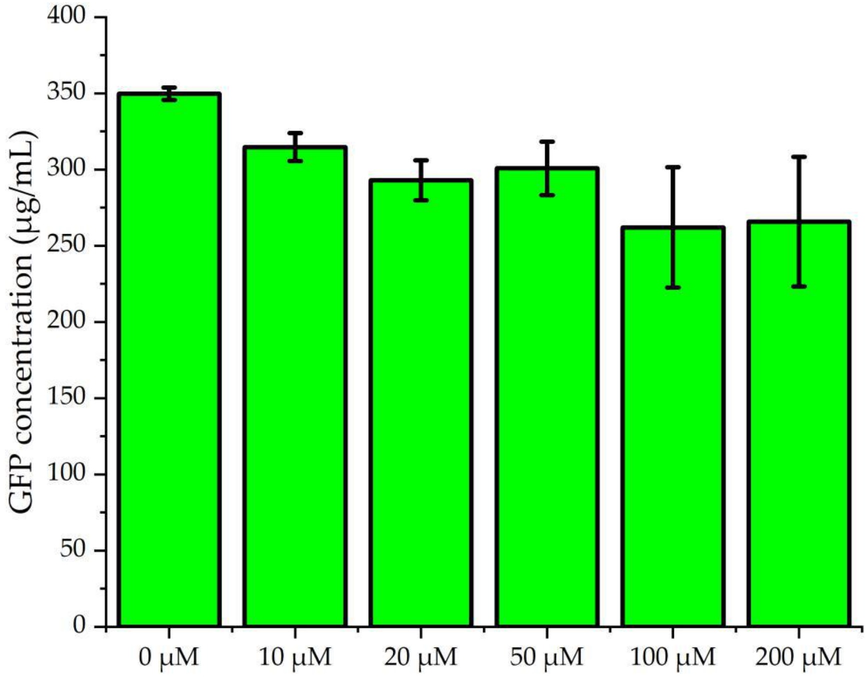


Figure S7: Endpoint measurements of cell-free *las* pathways treated by quercetin with varied concentrations.

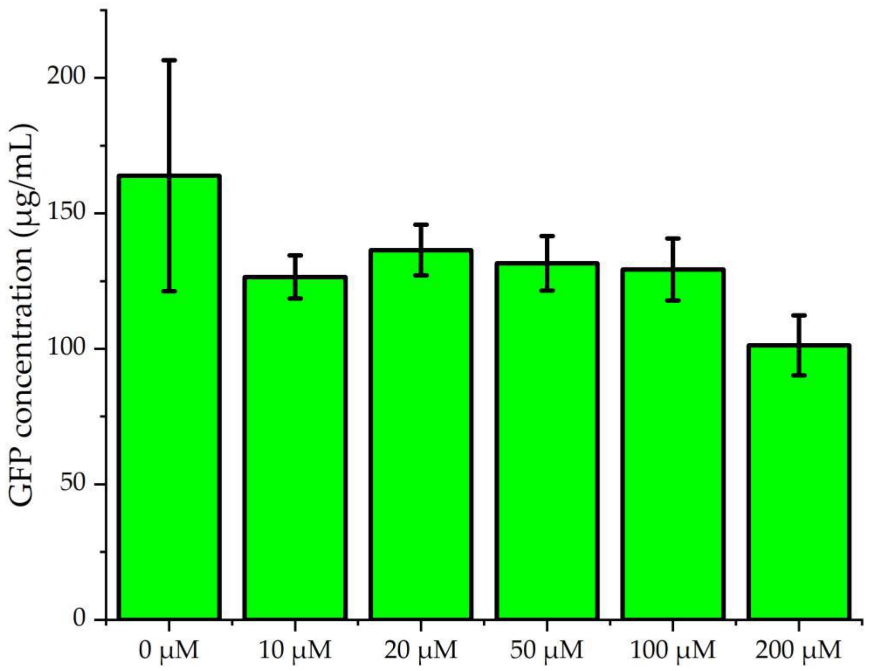


Figure S8: Endpoint measurements of cell-free reactions constitutively expressing GFP treated by quercetin with varied concentrations.
